# Potent Estrogen Receptor β Agonists with Inhibitory Activity *In Vitro*, Fail to Suppress Xenografts of Endocrine-Resistant Cyclin-dependent Kinase 4/6 inhibitor-Resistant Breast Cancer Cells

**DOI:** 10.1101/2024.01.12.575428

**Authors:** Lynn M. Marcho, Christopher C. Coss, Menglin Xu, Jharna Datta, Jasmine M. Manouchehri, Mathew A. Cherian

## Abstract

**Objective:** Seventy percent of newly diagnosed breast cancers are estrogen receptor-α positive and HER2/neu negative [1]. First-line treatments incorporate endocrine therapy and cyclin-dependent kinase 4/6 inhibitors [2]. However, therapy resistance occurs in most patients [3-5]. Hence, there is an urgent need for effective second-line treatments. We previously showed that the potent estrogen receptor-β agonists, OSU-ERb-12 and LY500307, synergized with the selective estrogen receptor modulator, tamoxifen, in vitro. Furthermore, we showed that these compounds inhibited endocrine-resistant and cyclin-dependent kinase 4/6-inhibitor-resistant estrogen receptor α-positive cell lines in vitro [6]. Here, we used fulvestrant- and abemaciclib-resistant T47D-derived cell line xenografts to determine the efficacy of the combination of OSU-ERb-12 and LY500307 with tamoxifen in vivo.

**Results:** Despite efficacy in vitro, treatments failed to reduce xenograft tumor volumes. Hence, we conclude that this treatment strategy lacks direct cancer cell-intrinsic cytotoxic efficacy in vivo.

## INTRODUCTION

Breast cancer is the most frequently diagnosed cancer in the world among women; estrogen receptor-α (ERα)-positive HER2-negative (ERα+/HER2-) breast cancer is the most common subtype [7, 8]. There are two nuclear estrogen receptors: ERα, which evidence has shown to be oncogenic in breast cancer, and estrogen receptor-β (ERβ), which is believed to be a tumor suppressor [9-14]. First-line treatments incorporate endocrine therapies, which suppress or block estrogen [15]. However, breast cancer cells inevitably develop resistance to endocrine therapy [16-18].

The development of cyclin-dependent kinase 4/6 (CDK4/6) inhibitors such as ribociclib, abemaciclib, and palbociclib, revolutionized the management of metastatic ERα+ breast cancer, doubling progression-free survival on first-line therapy [19-21]. However, these agents were introduced more than seven years ago, and many patients have since progressed [22]. Responses to second-line endocrine therapy are often brief. Hence, most of the ongoing ERα+/HER2-breast cancer research is focused on developing improved second-line therapies [2]. Our previous research indicated that highly specific agonists of ERβ inhibit the growth of ERα+ breast cancer cells *in vitro*, including endocrine-resistant and CDK4/6 inhibitor-resistant breast cancer cell lines. Moreover, our previous data suggested that selective ER modulators (SERM), such as tamoxifen, synergize with ERβ agonists against ERα+ breast cancer cell lines. In this report, we examined the potential efficacy of this strategy *in vivo* using a cell line xenograft model of endocrine and CDK4/6 inhibitor-resistant breast cancer.

## MATERIALS AND METHODS

### Drug formulation and drug administration

The Drug Development Institute (DDI) at The Ohio State University (OSU) synthesized OSU-ERb-012, as previously described [23]. OSU-ERb-12 was administered as a solution in 20% hydroxypropyl-β-cyclodextrin (HPBCD) (Sigma H107). Tamoxifen (Sigma T5648). LY500307 was synthesized and provided by the OSU DDI. Due to the hydrophobic nature of LY500307, extra light olive oil (Bertolli) was used as vehicle. Treatments included combination treatment with 100 mg/kg of OSU-ERb-012 + 20 mg/kg of tamoxifen in 100 μl of 20% HPBCD, given via oral gavage daily; 100 mg/kg of OSU-ERb-012 in 100 μl of 20% HPBCD, given via oral gavage daily; 20 mg/kg of tamoxifen in 100 μl of 20% HPBCD, given via oral gavage daily; 100 μl 20% of HPBCD, given via oral gavage daily; 10 mg/kg of OSU-ERb-012 + 20 mg/kg tamoxifen in 100 μl of 20% HPBCD, given via subcutaneous injection daily; 50 mg/kg of OSU-ERb-012 + 10 mg/kg of tamoxifen in 100 μl of 20% HPBCD, given via 2 subcutaneous injections in 2 separate locations on the body (for a total dose of 100 mg/kg of OSU-ERb-012 + 20 mg/kg of tamoxifen) daily; and 30 mg/kg of LY500307 + 20 mg/kg of tamoxifen in 100 μl of extra light olive oil given via oral gavage daily. Dose levels were chosen based on the pharmacokinetic properties of OSU-ERb-12 and were designed to provide peak plasma concentrations that exceeded the half maximal inhibitory concentration (IC_50_) of the drugs by at least 2-fold and to fully activate ERβ without activation of ERα [23].

### Cell culture

T47D cells (ATCC HTB-133; NCI-DTP Cat# T-47D, RRID : CVCL_0553), concurrently resistant to fulvestrant and abemaciclib, were selected by continuous culture in incremental concentrations over six months. Viability assays confirmed IC_50_ levels of more than 3 times that of the parental cell line, as described previously, thus verifying resistance to each agent [6]. Cells were diluted to a concentration of 5 million cells per 100 μl in phosphate-buffered saline (PBS) in individual syringes for mammary fat pad injection.

Cell pellets were prepared by allowing the fulvestrant- and abemaciclib-resistant T47D cell line to grow overnight, followed by treatment with 5 μM, which was the estimated IC50, of OSU-ERb-12 or LY500307 for 72 hours. Control cells were treated with dimethyl sulfoxide (DMSO) vehicle. Fresh media and drugs were replaced every alternate day. After 72 hours of treatment, the cells were collected and centrifuged to obtain a cell pellet.

### Mouse maintenance

Animal care, housing, and experimental procedures were under the ordinance of The Ohio State University’s Institutional Animal Care and Use Committee (IACUC), protocol number 2019A00000040. Animals were maintained on a 12-hour light/12-hour dark schedule and had free access to standard food and water. Ovariectomized NCG mice (strain 572, NOD-SCID-γ^-/-^) that are known to lack T cells, B cells, and NK cells (Charles River Laboratory) were used for cell-line xenografts of T47D cells. Mice were euthanized by carbon dioxide inhalation and confirmed with cervical dislocation.

### Subcutaneous implantation of estradiol pellet

From day -3 to day 5, the mice received 0.2 mg of ibuprofen/ml of drinking water. On day -2, the mice were anesthetized and received a 0.1 mg/kg subcutaneous injection of buprenorphine. Hair was shaved from the upper back between the shoulder blades. The exposed skin was cleaned with alternating rounds of betadine and 70% ethanol and repeated three times each. Press’n Seal (GLAD) was used as a surgical drape over the surgery area. A 10G trocar (Innovative Research of America, MP-182) was used to implant the 60-day release 0.72 mg 17β estradiol pellet (Innovative Research of America, SE-121) under the skin. The wound was closed with a drop of Vetbond Tissue Adhesive. Mice recovered under a heat lamp until ready to return to their home cages.

### Mouse mammary fat pad injection

On day 0, the mice were anesthetized and received a 1 mg/kg subcutaneous injection of buprenorphine. The mice continued to receive 0.2 mg of ibuprofen/ml of drinking water. The mice were shaved in the inguinal region to access the lower mammary fat pad. The skin was cleaned with betadine and 70% ethanol, and a drape was used as described above. A 1 cm-long longitudinal incision was made, and the mammary fat pad was identified. Cells were injected directly into the fat pad, which was then tucked back inside the incision. The wound was closed with a drop of tissue adhesive.

### Data collection

Mice were weighed before drug treatments began and twice weekly during the experiment. After tumor cell injection, the length and width of the tumor were measured twice weekly using calipers. The tumor volume was calculated using the following formula: (4/3) x π*L x W x W, where width was the smaller of the two length measurements. Once the tumor reached a volume of 500 mm^3^, treatment began. Treatment groups were randomly assigned to mice via a random group generator. Each treatment group included 8-9 mice. Treatments were administered 5 days a week for 6 weeks for a total of 30 doses. Twice weekly tumor measurements continued during treatment administration. Study termination criteria included a tumor volume measuring 1.6 cm in diameter, tumor ulceration, or morbid condition. Mice were euthanized after 6 weeks of treatment, and tumors were removed, flash-frozen in dry ice, and stored at -80°C.

### Blood collection and drug concentration analysis

Blood was collected via submandibular vein sampling 2 hours and 24 hours after drug administration. The drug concentrations of OSU-ERb-12 and LY500307 in cell pellets, plasma, and tumors were measured by Charles River Laboratories (Wilmington, MA) by high-performance liquid chromatography/mass spectrometry. Drug concentrations were expressed in micromoles/L.

### Statistical analysis

Group differences in tumor volume changes over time were analyzed with a linear mixed model using R 4.10 (Vienna, Austria). In the linear mixed model, time points, treatment groups (with vehicle treatment as the reference), and interactions between time and treatment groups served as predictors. Group differences in tumor weight were assessed with a one-way analysis of variance (ANOVA) model, and partial *η*^*2*^ was used to indicate effect size. Group differences in mouse survival status (Y/N) were evaluated with a logistic regression model. Group differences in LY500307 drug concentration were assessed using Wilcoxon’s rank sum test. SPSS version 28.0 (IBM Corp., Armonk, NY) was used for these calculations. Group differences in mouse weight changes over time were analyzed by linear mixed model using R 4.10. For this model, time and treatment groups (vehicle as the reference) and interactions between treatment and time served as predictors. Pairwise comparisons of OSU-ERb-012 drug concentration were assessed using Dunn’s test with concentration in the cell pellet serving as the reference group. GraphPad Prism version 10.0.2 software (Boston, MA) was used to calculate this test, and a two-sided alpha level of 0.05 was adopted.

## RESULTS

### Tumor volume and weight

Tumor volume, measured twice weekly, increased over time across all treatments (*p*≤0.001) (**Figure 1A**). However, there was no statistically significant difference between any treatment group and the vehicle-treated group over 6 weeks of drug administration. Tumors were harvested after completion of 6 weeks of drug administration. There was no significant difference between tumor weight of any treatment group and vehicle treated group (*p*=0.969) (**Figure 1B**). All treatment groups demonstrated a small magnitude of effect on tumor weight as indicated by partial ⍰^2^ = 0.029.

**Figure 1:**
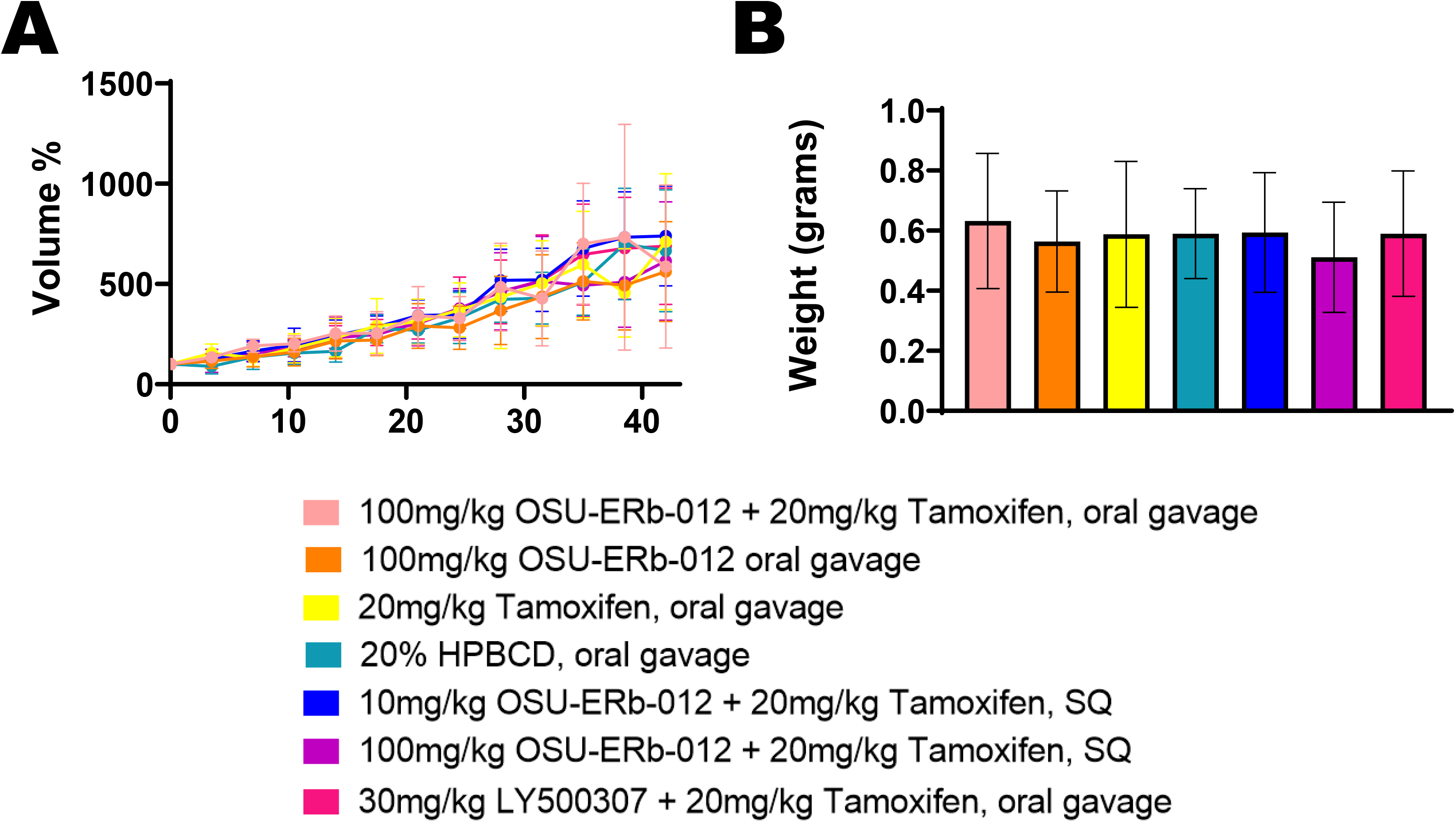
Tumor Volume and Tumor Weight. **(A)** Depicted tumor volumes were normalized to their pretreatment volumes (pretreatment volumes set at 100%). The average tumor volume per treatment group is shown by the date of treatment. A linear mixed model showed that tumor volume significantly increases with time. Beta coefficient = 324.096, standard error = 19.743, *p*≤0.001. None of the treatments were significantly different compared to vehicle control. **(B)** The average tumor weight per treatment group after completion of 6 weeks of drug administration is shown. One-way ANOVA results showed that there was no significant difference in tumor weight between each treatment and vehicle treatment. ***p***=0.969, partial ⍰^2^ = 0.029, indicating a small magnitude of effect size. *(*, p<0*.*05)*.

### Mouse survival and weight

Mouse weight decreased over time, indicating a possible adverse or toxic effect of the treatments or tumor burden (**Figure 2A**). Compared to vehicle treatment, the OSU-ERb-12 oral gavage (*p*=0.001) and 100 mg OSU-ERb-12 + tamoxifen subcutaneous (SQ) (*p*<0.001) groups lost significantly more weight over time, while the group treated with tamoxifen alone (*p*<0.001) lost significantly less weight over time.

**Figure 2:**
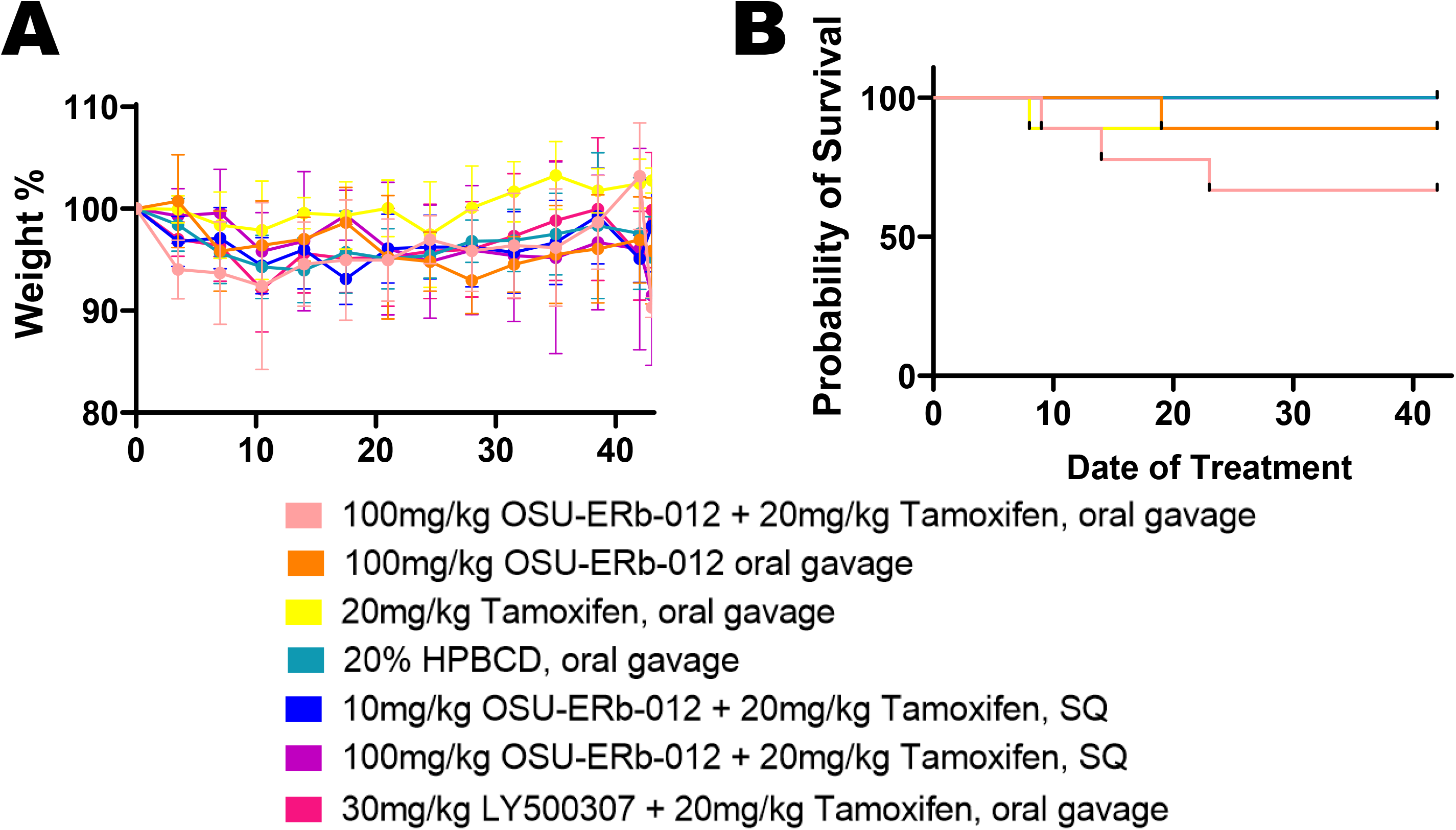
Mouse Weight and Mouse Survival. **(A)** Mouse weights were normalized to begin at 100% at the beginning of treatment. Linear mixed model results showed that overall mouse weight decreased significantly over time. With the 20% HPBCD vehicle treatment being the reference group, the OSU-ERb-12 oral gavage treatment (*p*=0.001) and 100 mg OSU-ERb-12 + tamoxifen SQ treatment (*p*<0.001) significantly enhanced the decreasing trend, while the tamoxifen alone group (*p*<0.001) significantly weakened the decreasing trend. **(B)** Mouse Survival: Logistic regression showed that none of the treatment groups were significantly different from vehicle treatment in terms of mortality. *(*, p<0*.*05)*.

Two mice on OSU-ERb-12 + tamoxifen oral gavage treatment expired during treatment; a third was euthanized due to a veterinary concern. One mouse on OSU-ERb-12 oral gavage treatment expired during treatment. One mouse each from the tamoxifen alone and LY500307 + tamoxifen treatment groups was euthanized due to veterinary concerns. None of the treatment groups were significantly different from the vehicle treatment group in terms of survival (**Figure 2B**).

### Drug level concentrations in plasma and tumors

The concentration of OSU-ERb-012 in the cell pellet was 290-fold higher (*p*=0.002, pellet mean rank = 22.50, tumor mean rank = 10.50) compared to the tumors (**Figure 3A**). The peak concentration of LY500307 in the plasma was a little more than half of the IC50 that was established in vitro for endocrine-resistant and CDK4/6 inhibitor-resistant T47D cells [6] (**Figure 3B**). The concentration of LY50037 in the cell pellet was 1,076-fold higher than in the tumors (*p*=0.021, pellet mean rank = 6.50, tumor mean rank = 2.50) (**Figure 3C**).

**Figure 3:**
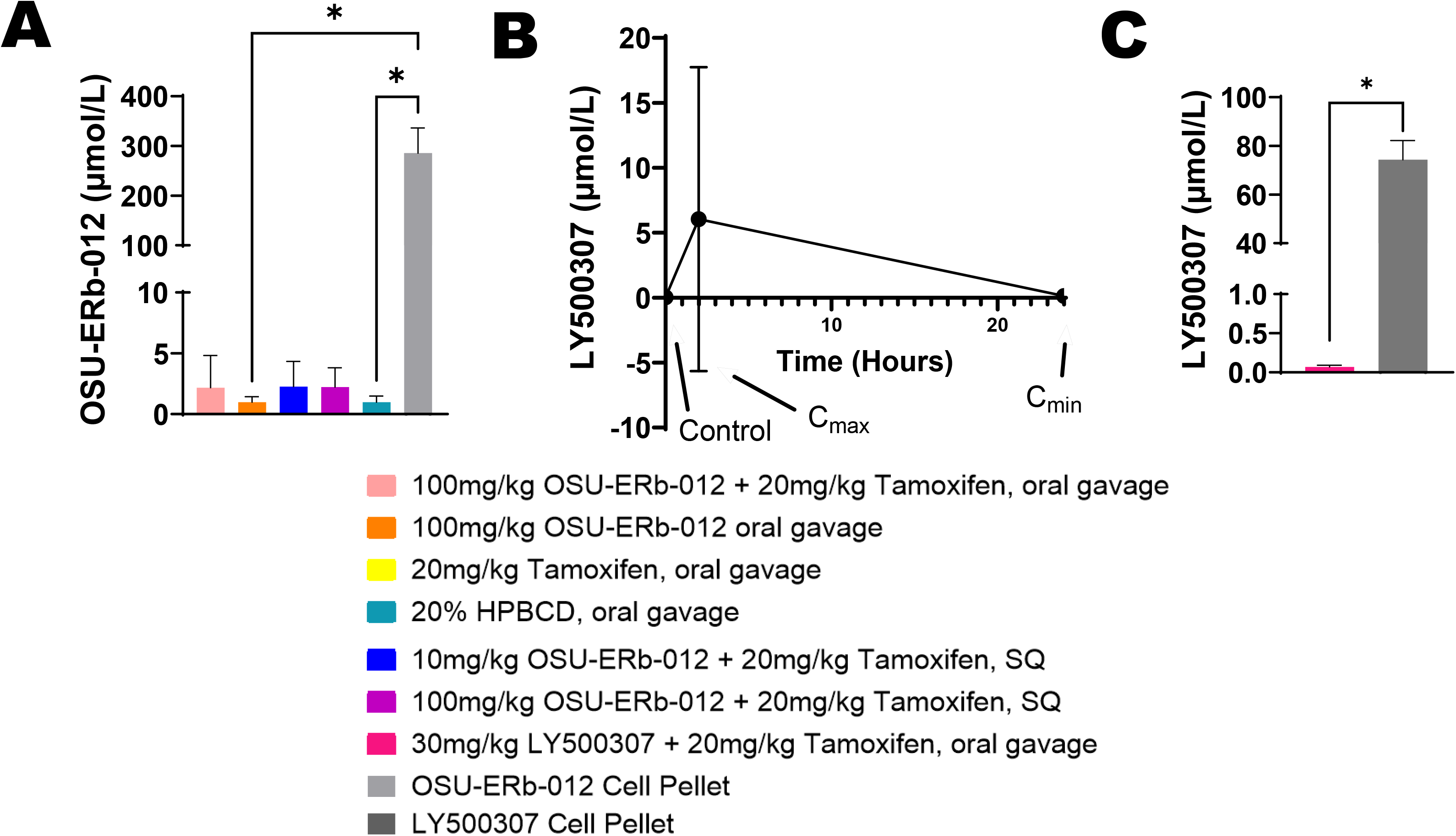
Drug Level Concentrations. **(A)** OSU-ERb-012 concentration in tumors compared to cell pellets. Pairwise comparisons using Dunn’s test showed that the concentration of OSU-ERb-12 in the OSU-ERb-12 treated cell pellets was significantly higher than OSU-ERb-12 oral gavage treated tumors (*p*=0.0320) and 20% HPBCD vehicle-treated tumors (*p*=0.0465). Drug concentrations in other groups were not statistically significantly different from those in OSU-ERb-12-treated cell pellets, despite a 100-fold difference, due to multiple comparisons. **(B)** LY500307 concentration in plasma of mice treated with LY500307. (C) LY500307 concentration in tumors compared to cell pellets treated with LY500307. The Wilcoxon rank sum test showed that the concentration of LY500307 in cell pellets was significantly higher than that in LY500307 treated tumors (*p*=0.021). *(*, p<0*.*05)*.

## DISCUSSION

The 100 mg/kg treatment dose of OSU-ERb-12 was 10-fold higher than pharmacokinetic testing suggested was necessary to achieve peak plasma concentrations (unpublished data). This suggests that peak concentrations greater than the IC_50_ could be achieved with this dosing schedule. Despite being dosed aggressively, the drug combinations lacked efficacy *in vivo*, which was unexpected based on our in vitro results [6]. As a comparison, *in vitro* cells were also treated with the ERb agonists to determine what cellular concentration were possible, in the absence of the complex systemic environment.

We believe that the lack of efficacy may be due to the low intratumoral drug concentrations, which were significantly less than those achieved in pellets of cells treated *in vitro* at the IC_50_ concentration. The concentration of OSU-ERb-012 in cell pellets was 290-fold higher compared to intratumoral concentrations, and the concentration of LY500307 in cell pellets was 1076-fold higher than intratumoral concentrations. The concentration of LY500307 in plasma was similar to that in tumors.

Given the hydrophobic nature of the ERβ agonists tested, drug transport or plasma protein binding may have resulted in low “free” concentrations. Additionally, we hypothesize that the lipophilicity of OSU-ERb-12 may have led to its concentration in lipid-rich subcellular structures. This would offer an explanation for why the drug concentrations in cell pellets were so much higher than the intratumoral concentrations. Hence, although we utilized a dose that was 10-fold higher than pharmacokinetic testing suggested was necessary, we failed to achieve intratumoral concentrations that were established to be cytotoxic *in vitro*.

## LIMITATIONS

One limitation of this study is the lack of a pharmacodynamic marker for ERβ activation. However, unique markers for ERβ-specific activation are not well established [24].

## ABBREVIATIONS

ANOVA: analysis of variance
CDK4/6: cyclin-dependent kinase 4/6
DMSO: dimethyl sulfoxide
EC_50_: half maximal effective concentration
ERα: estrogen receptor-α
ERβ: estrogen receptor-β
HPBCD: hydroxypropyl β cyclodextrin
IC_50_: Half maximal inhibitory concentration
PBS: phosphate buffered saline
SERM: selective ER modulators
SQ: subcutaneous
μl: microliter
μM: micromolar
mg: milligram
kg: kilogram

## DECLARATIONS

### Ethics approval and consent to participate

All methods were carried out in accordance with IACUC’s regulations and guidelines.

### Consent for publication

Not applicable

### Experimental procedures and protocols

Animal care, housing, and experimental procedures were under the ordinance of The Ohio State University’s Institutional Animal Care and Use Committee (IACUC), protocol number 2019A00000040. Additionally, all methods were carried out in accordance with IACUC’s regulations and guidelines.

### Availability of data and material

The datasets used and/or analyzed during the current study are available from the corresponding author upon reasonable request.

### Author’s contributions

MAC conceived of this project. LMM, CCC, and MAC designed experiments. LMM performed most of the experiments and JD prepared the cell pellets. LMM and MX performed statistical analysis. LMM authored the draft. JMM and MAC edited the manuscript.

## Acknowledgments

The authors would like to thank scientific editor Angela Dahlberg, Division of Medical Oncology, The Ohio State University Comprehensive Cancer Center, for editing this manuscript.

